# Non-target impacts of fungicide disturbance on phyllosphere yeasts in different crop species and management systems

**DOI:** 10.1101/2021.09.20.461135

**Authors:** Zachary A. Noel, Reid Longley, Gian Maria Niccolò Benucci, Frances Trail, Martin I. Chilvers, Gregory Bonito

## Abstract

- Fungicides reduce fungal pathogen populations and are essential to food security. Fungicide disturbance of plant microbiomes has received limited attention. Understanding the impacts of fungicides on crop microbiomes in different cropping systems is vital to minimizing unintended consequences while maintaining their use for plant protection.
- We used amplicon sequencing of fungi and prokaryotes in maize and soybean microbiomes before and after foliar fungicide application in leaves and roots from plots under long-term no-till and conventional tillage managements. We examine fungicide disturbance and microbiome resilience across these treatments.
- Foliar fungicides directly affected phyllosphere fungal communities, but not root fungal communities or prokaryote communities. Impacts on fungal phyllosphere composition and resiliency were management-dependent and lasted more than thirty days. Fungicides lowered pathogen abundance in maize and soybean and decreased the abundance of Tremellomycetes yeasts, especially the Bulleribacidiaceae, including core microbiome members.
- Fungicide application reduced network complexity in the soybean phyllosphere. Bulleribacidiaceae often co-occurred with *Sphingomonas* and *Hymenobacter* in control plots, but co-occurrences were altered in fungicide plots. Results indicate that foliar fungicides lower pathogen and non-target fungal abundance and may impact prokaryotes indirectly. No-till management was more resilient following fungicide disturbance and recovery.

## Introduction

Disturbances from chemical applications in agriculture reduce the abundance of pests and pathogens and are common in modern agricultural ecosystems (Rykiel, 1985; Glasby & Underwood, 1996; Sullivan & Sullivan, 2003; Landers *et al*., 2012; Shade *et al*., 2012). However, applying disturbance concepts to microbial communities can be challenging to assess recovery and analyze the full impacts. A lack of data on the impacts crop management combined with fungicide disturbances on the plant microbiome hinders developing novel strategies to minimize diversity loss, understand unintended consequences of these applications, and improve crop microbiomes’ resilience. Observing fluctuations in taxa abundance and secondary effects mediated through microbial interactions following fungicide application opens the possibility for novel ecologically motivated strategies that promote microbiome stability or resilience following a fungicide application.

Fungicide use has become common in conventional agricultural systems. Yet, concerns remain about direct and indirect effects on non-targeted organisms, consequences (i.e., resistance), and negative impacts on the environment or human health (Hahn, 2014; Schaeffer *et al*., 2017b; Zubrod *et al*., 2019). The rapid evolution of fungicide resistance in plant and human pathogenic fungal populations can cause devastating epidemics in agricultural ecosystems, with spill-over effects to public health (Verweij *et al*., 2009; Delmas *et al*., 2017; Riat *et al*., 2018; McDonald *et al*., 2019). For example, there is substantial concern about the overuse of azole fungicides that have been linked to the resistance of *Aspergillus fumigatus* to antifungals in human infections (Riat et al. 2018; Verweij et al. 2009). Despite concerns, foliar fungicide applications in maize (*Zea mays* L.) and soybean (*Glycine max* L. Merr) are often made without substantial pathogen pressure due to perceived or marketed yield benefits (Wise & Mueller, 2011; Kandel *et al*. 2021). A meta-analysis of soybeans demonstrated that foliar fungicide application in the absence of disease increased yield by 2.7%. Still, economic analyses indicated that fungicides are less profitable without disease pressure (Kandel *et al*., 2021). While fungicides are necessary for crop protection, minimizing non-target effects and unintended consequences is critical in evaluating the sustainability of agricultural production systems.

Studies reporting fungicidal and pesticidal impacts to soil microbiomes (Imfeld & Vuilleumier, 2012; Fournier *et al*., 2020) and aquatic organisms (Zubrod *et al*., 2019) are more abundant than those where fungicides are applied directly to the foliage, which is surprising considering the numerous phyllosphere pathogens and microorganisms present in this habitat. From the few studies focused on the plant phyllosphere, a consistent non-target effect is detected against phyllosphere yeasts. For example, one study on grapevine microbiomes reported minimal and transient impacts to the phyllosphere microbiome, including phyllosphere yeast abundance (Perazzolli *et al*., 2014). Similarly, repeated application of broad-spectrum fungicides has been shown through culture-based and culture-independent methods to decrease phyllosphere yeast richness (Dickinson & Wallace, 1976; Southwell *et al*., 1999; Sapkota *et al*., 2015; Knorr *et al*., 2019).

Yeasts that inhabit the phyllosphere are well suited to oligotrophic and dynamic environmental conditions present on leaf surfaces and have been consequently used as biocontrol of plant pathogens (Freimoser *et al*., 2019). They are known to produce extracellular polysaccharides and surfactants, which may be necessary for creating or maintaining biofilms (Fonseca & Inácio, 2006). Additionally, some phyllosphere yeasts, including species of basidiomycete yeasts in *Cryptococcus* and *Sporidiobolus*, produce carotenoid compounds, which have antioxidant properties and may protect the yeasts and potentially other resident microbes from stress in the phyllosphere (Cobban *et al*., 2016). Phyllosphere yeast communities have also been linked to pollinator insects by altering floral nectary chemistry, and fungicides can modify this relationship (Cadez *et al*., 2010; Schaeffer *et al*., 2017a). However, no study has addressed the links between phyllosphere yeasts and other phyllosphere residing microorganisms. While indirect and collective effects of removing single species or groups of species from ecosystems proposed in ecological theory since the 1940s and have been studied in various macro-organism contexts such as conservation biology, disturbance ecology, and food web ecology, such effects are comparatively understudied in microbiomes (Tilman, 1999; Ripple & Beschta, 2005; Sahasrabudhe & Motter, 2011). In microbiomes, network complexity (i.e., linkage density) has been correlated to ecosystem functioning and stability (Zhou *et al*., 2010; Wagg *et al*., 2019). Consequently, indirect effects may be revealed through co-occurrence patterns, which may not otherwise be seen through culturing or sequencing one target group.

Since the US Dust Bowl of the 1930s, soil conservation efforts have led to the steady adoption of minimum or no-till agriculture management systems (Claassen *et al*., 2018). Cropping management systems have been demonstrated to impact phyllosphere microbiomes (Gdanetz & Trail, 2017; Longley *et al*., 2020). Crop management’s effect on the resilience of foliar fungal communities following fungicides has not been explored, but differing impacts of fungicides in different agricultural managements are probable. In the only study of this kind, agricultural management altered the response of microbial communities to tetraconazole fungicide application (Sułowicz *et al*., 2016). However, the two management treatments existed in different locations confounding efforts to distinguish fungicide management response effects from those of location (Karlsson *et al*., 2014). Long-term experiments such as the Main Cropping Systems Experiment at the Kellogg Biological Research Station, circumvent these confounding effects by applying all treatments at a single location.

Here, we characterize non-target effects of foliar fungicides and resilience of the maize and soybean leaf and root microbiomes in no-till and conventional plots of the Long-term Ecological Research (LTER) Main Cropping Systems Experiment at the Kellogg Biological Station (KBS). Our research objectives were three-fold: (1) to determine whether fungicides alter microbial diversity across plant compartments (e.g., leaves or roots), crop species (e.g., maize or soybean), or management (conventional vs. no-till); (2) to identify non-target and indirect effects of fungicide applications, and (3) determine if crop management alters the resiliency of the microbiome. We hypothesized that fungicides would alter both maize and soybean microbial (fungal and prokaryotic) diversity and network complexity. We predicted that this effect would be pronounced in the leaves compared to roots. Additionally, given that plant microbiomes have been shown to differ under the two management systems (Longley *et al*., 2020), we hypothesize that the response and recovery of plant microbiomes following fungicides would also differ between the two management treatments. This LTER site allows for a novel approach by eliminating any differences caused by location bias and assessing the effect of fungicide application under long-term agricultural management. We apply a novel microbiome network analysis approach to determine the impact fungicides have on prokaryote-fungal co-occurrences in the plant microbiome. Finally, we used random forest models to predict prokaryote taxa responsive to altered fungal diversity demonstrating the possible indirect effects of fungicides.

## Materials and Methods

### Experimental design

#### Sample Site and Management Systems

All samples were collected from the no-till and conventional management systems of maize (*Zea mays* L.), soybean (*Glycine max* L. Merr), and wheat (*Triticum aestivum* L.) rotation of the main cropping experiment at Michigan State University’s Kellogg Biological Station (KBS) Long Term Ecological Research (LTER) site. The site contains one-hectare plots, consistently managed since 1989 (Robertson & Hamilton, 2015). Each agricultural management at the site is represented in six replicate plots spread randomly throughout to avoid location bias. Fungicide micro-plots (10 feet wide x 20 feet long) were established within four of the six replicate plots under no-till and conventional management treatments. Control samples were taken from micro-plots of the same size directly next to the fungicide micro-plots. Samples were taken from the middle of plots or towards the edge of the microplot that was not along the fungicide-control (i.e., at least 6 meters apart) border to minimize the effect of spray drift. In 2017, the fungicide Headline® was applied to maize foliage at a recommended rate of 877 ml ha^-1^ (12 fl oz acre^-1^). Headline® contains the active ingredient pyraclostrobin, which acts as a mitochondrial respiration inhibitor. Pyraclostrobin is a local penetrant fungicide with translaminar movement and is not translocated in the xylem (Latin, 2017). In 2018, soybean foliage was sprayed with Delaro® fungicide on 3 August 2018 (Bayer, Raleigh, NC, USA) at a recommended rate of 731 ml ha^-1^ (11 fl oz acre^-1^). The active ingredients in Delaro® are a combination of trifloxystrobin which inhibits mitochondrial respiration, and prothioconazole, which inhibits ergosterol synthesis. Trifloxystrobin is a local penetrant fungicide with translaminar movement and is not translocated in the xylem (Latin, 2017). Prothioconazole has acropetal penetrant activity and has weak basipetal movement (Augusto & Brenneman, 2012).

#### Sample collection and DNA extraction

In the summer of 2017, maize leaf and root samples were collected at three time points throughout the growing season. The first sampling occurred before the fungicide application on 26 June 2017 (V6 growth stage), the second was 9-days post fungicide application (V8 growth stage), and the final sampling was 35-days post fungicide application (V15 growth stage). Leaves and roots from three plants from four replicate control or adjacent fungicide treated plots of each no-till, and conventional management was sampled at each time point. Maize leaves were sampled by removing two whole leaves from each plant and placing them into a sterile Whirl Pak (Nasco, Madison, WI, USA) for transport back to the lab where they were stored at -80ºC until they were lyophilized. At the V6 and V8 growth stage, the sixth and seventh leaf was sampled. However, at the V15 growth stage, three leaves above the ear leaf were sampled. Roots were sampled by removing whole plants from the soil and the entire root system to the soil line. Then roots were washed in the field before being transported back to the lab, where roots were washed again with 0.1% tween 20 (ThermoFisher Scientific, USA) and deionized water. Samples were stored at -80ºC before being lyophilized for DNA extraction. Following lyophilization, the fine roots were removed from the root system and used for DNA extraction.

In 2018, soybean leaves were sampled at three time points during the growing season, with the first occurred before fungicide spray on 3 August 2018 (R3 growth stage), the second occurred 13-days post fungicide application (R4 growth stage), and the final occurring 33-days post fungicide application (R6 growth stage) (Fehr *et al*., 1971). Soybean leaves were sampled with a flamed metal hole punch, washed in 80% ethanol, and flame sterilized between samples. Three 6-mm leaf discs from three separate leaves were punched directly into an Eppendorf tube (Eppendorf, Germany) containing 500 μl of CSPL buffer (Omega Bio-Tek, Norcross, GA, USA). As with the maize roots, soybean roots were removed at the soil line and placed into a new Whirl-Pak (Nasco, Madison, WI, USA) bag containing approximately 50 ml of 0.1% tween 20 to remove the remaining soil. Root samples were transported back to the lab, where roots were washed again with DI water, and samples were stored at -80ºC until processing. Maize and soybean leaf and root tissue were pulverized for 2-min at a speed of 30 Hz with two 4-mm stainless balls in a TissueLyser II (Qiagen, Venlo, Netherlands). Total DNA was extracted from plant tissues with the OMEGA Mag-Bind Plant DNA Plus kit (Omega Bio-Tek, Norcross, GA, USA) following the manufacturer’s instructions with the aid of a KingFisher Flex™ liquid handling machine (ThermoFisher Scientific, USA). Five or six internal negative extraction controls were included per 96-well plate in each DNA extraction.

#### Amplicon library preparation for ITS and 16s community profiling

Amplicon libraries were prepared from a three-step PCR protocol as modified from Lundberg et al. (2013) and described by Longley et al. (2020). In brief, fungal libraries were constructed around the ITS and were amplified using the primers ITS1F and ITS4 (Gardes & Bruns, 1993). Prokaryote libraries targeted the V4 region of 16S rRNA with the primers 515F and 806R (Caporaso *et al*., 2011). Supporting information Tables S1, S2 and S3 describe PCR protocols, primers, and cycling conditions in detail. Amplicons were purified with the SequalPrepTM Normalization Plate Kit (ThermoFisher Scientific, USA) and then pooled and concentrated with Amicon® Ultra 0.5 mL filters (EMDmillipore, Germany).

Subsequently, the library was purified, and size selected with Agencourt AMPure XP magnetic beads (Beckman Coulter, USA). Amplicon libraries were quantified and checked on the Agilent 4200 TapeStation DNA10000 and Kapa Illumina Library Quantification qPCR assays. All amplicon libraries were then paired-end sequenced (300 bp reads) on an Illumina MiSeq with a v3 600 cycles kit (Illumina, USA).

#### Bioinformatic sequence processing

Fungal ITS1 or prokaryotic 16S V4 reads were demultiplexed in QIIME 1.9.1 (Caporaso *et al*., 2010). Forward and reverse prokaryote reads were merged using QIIME 1.9.1. Only forward fungal ITS1 reads were used since reverse reads did not overlap. After removing primers with Cutadapt 1.8.1 (Martin, 2011), fungal reads were trimmed to remove the conserved SSU and 28S regions. Reads were then quality filtered at an expected error threshold of 0.1 and truncated to equal length (fungi 200 bp; prokaryote 300 bp) in USEARCH 11.0.667 (Edgar, 2010). Singletons and chimeras were removed, and *de novo* OTU clustering was performed at a 97% similarity using the UPARSE algorithm (Edgar, 2013). Using CONSTAX2 (Gdanetz *et al*., 2017; Liber *et al*., 2021), the taxonomic classification of fungal and prokaryotic OTU’s representative sequences was performed against the UNITE eukaryote database, ver. 8.2 of 04.02.2020 (Abarenkov *et al*., 2020) and SILVA, version 138 (Quast *et al*., 2013), respectively. To filter out non-target taxa and OTUs unidentified at the Kingdom level, CONSTAX was run twice under different cutoff levels, as suggested by Bowsher *et al*. (2020). Non-target taxa, OTUs not assigned to a Kingdom, and OTUs identified as either chloroplast or mitochondria in either database were removed from further analysis (Zhang *et al*., 2019).

#### Statistical analysis

Data were imported into R 4.0.3 (R core team 2020), and the R packages *phyloseq* 1.24.2 (McMurdie & Holmes, 2013) and *vegan* 2.5.3 (Jari Oksanen *et al*., 2018) were used for most analyses. Samples with low sequencing coverage (less than 1000 reads) were removed from the analysis. Contaminant OTUs (i.e., those prevalent in negative extraction controls) were removed with the R package *decontam* (Davis *et al*., 2018). Before normalization, richness was assessed for Prokaryotes and Fungi in the leaves and roots of each crop using the ‘estimate_richness’ function in the *phyloseq* package. Results of alpha diversity analyses were plotted using the ‘ggplot2’ package (Wickham, 2009). Then, sample read counts were normalized using the cumulative sum scaling technique within the *metagenomeSeq* R package (Paulson *et al*., 2013). Differences in fungal and prokaryotic community centroids were tested through permutational multivariate analysis of variance (PERMANOVA) with the ‘adonis2’ function. Variation in multivariate dispersion was tested with the ‘betadisper’ function. More specific hypotheses were tested based on constrained analysis of principal coordinates (CAP) (Anderson & Willis, 2003).

Differentially abundant taxa resulting from fungicide application were identified by comparing fungicide treated plots to control plots through an analysis of the composition of microbiomes (ANCOM v 2.1) (Mandal *et al*., 2015). For differential abundance analysis, fungal OTUs (fOTU, hereafter) with a mean relative abundance less than 1.0^-5^ and fOTUs with zeros present in 95% samples were discarded from the analysis to avoid detecting fOTUs as significantly different based on stochasticity. Additionally, fOTUs that were never present in fungicide treated plots were not included. Fungal OTUs were determined to be significant if the W value was greater than 70% of the taxa tested based on Wilcoxon ranked sum test between additive log-ratio transformed data and a Benjamini-Hochbergj adjusted P-value (α = 0.05) (Mandal *et al*., 2015). Recovered taxa (i.e., transient effects) were defined as fOTUs that were significantly less abundant following fungicide treatment but were not significantly less abundant from non-disturbed plots, after 33- or 34-dpf, for soybean or maize, respectively. Non-recovered taxa were defined as those fOTUs with significantly altered abundance following fungicide application and remained significantly altered for the remainder of the sampling. Additionally, a portion of non-recovered taxa was also locally extinct, which were defined as taxa present before fungicide application but having zero relative abundance following fungicide application in fungicide treated plots through the remainder of the sampling while being present in the control plots. Finally, taxa that did not have significantly altered abundance following fungicide application but then had significantly different abundance at a later sampling point (i.e., 33- or 34-dpf) were defined as indirect effects.

The core phyllosphere fungal and prokaryotic taxa from the non-fungicide disturbed no-till or conventional plots were identified based on each abundance and occupancy across space and time. Taxa that contributed to the last 2% increase in Bray-Curtis distances were defined as the core (Shade & Stopnisek, 2019).

We built random forest regression models to test the effect of altered prokaryote abundance through fungal diversity by using prokaryote abundances to predict fungal diversity. Random forest models were generated with the ‘randomForest’ function in the *randomForest* R package (Liaw & Wiener, 2002). To remove redundant features and avoid overfitting models, we removed redundant OTUs with the ‘Boruta’ function in the package *Boruta* (Kursa & Rudnicki, 2010). The method performs a top-down search for relevant OTUs by comparing the importance of the original OTUs from those selected at random. Models were tuned to achieve the lowest stable out-of-bag (OOB) error estimate possible, and the best *mtry* value (number of OTUs sampled at random in the entire pool for each tree at each split) was selected using the ‘tuneRF’ function in *randomForest* R package.

Network analysis was conducted on soybean and maize leaf samples to estimate co-occurrences among prokaryotic and fungal OTUs in each host and determine whether fungicides altered fungal-prokaryotic co-occurrences and network complexity (i.e., linkage density). For network analyses, soybean and maize fungal and prokaryotic OTU tables were filtered to exclude taxa with mean relative abundance below 1.0^-5^. A co-occurrence meta-matrix was estimated using the Meinshausen and Bühlmann algorithm within the *SpiecEasi* R package with the ‘nlambda’ set to 100 and with ‘lambda.min.ratio’ set to 1.0^-2^ (Kurtz *et al*., 2015). From this meta-matrix, subnetworks were created from taxa present within each sample. Then, network complexity was calculated for each subnetwork. The contribution of the Bulleribacidiaceae to network complexity was assessed by examining the change in the cumulative edge weights across subnetworks with prokaryotic genera.

## Results

### General sequencing results

The final fungal OTU table contained 20,844,912 ITS1 reads across 554 samples, including 5315 fOTUs after filtering 36 contaminant fOTUs detected in negative controls. The median read depth was 30,370 ITS1 reads per sample. Prokaryotes contributed 29,691,681 total reads across 555 samples with a median read depth of 47,590 reads per sample. A total of 14,291 prokaryote OTUs (pOTU, hereafter) were defined after filtering 55 contaminant pOTUs detected in the negative controls. Rarefaction curves verified that the median read depth adequately sampled the diversity present (Fig. S1).

### Fungicides alter the maize and soybean leaf fungal richness

Following fungicide application, the richness of maize and soybean leaf fungal communities was significantly reduced (P < 0.05) compared to that of non-sprayed control plots (Fig. S2). This reduction in fungal richness was observed in both crops in each management except no-till maize samples. This effect was most pronounced for the fungal classes of Dothideomycetes (target) and Tremellomycetes (non-target). However, in other fungal classes such as the Sordariomycetes, there was no significant difference in richness between control and fungicide treated samples following fungicide applications. Therefore, reductions in fungal richness were likely confined to specific fungal classes instead of being across all classes. Among prokaryotes, there were no consistent differences between control and fungicide samples in either crop. Likewise, there was no evidence that fungicides altered fungal or prokaryotic richness in roots.

### Fungicides alter the maize and soybean leaf fungal community composition

Fungal and prokaryote community composition varied significantly through time (i.e., days post fungicide or dpf) and crop management in maize and soybean leaves and roots (Table S4; Table S5). Notably, before fungicides were sprayed, crop management was shown to have a significant effect on the maize and soybean phyllosphere fungal and prokaryotic communities (maize leaf fungi R^2^ = 0.050, *P* = 0.001; maize leaf prokaryotes R^2^ = 0.038, *P* = 0.005; soybean leaf fungi R^2^ = 0.058, *P* = 0.020; soybean leaf prokaryotes R^2^ = 0.046, *P* = 0.049). Furthermore, fungal and prokaryotic phyllosphere community compositions in control and treatment plots were indistinguishable from each other prior to applying fungicide treatments (maize leaf fungi R^2^ = 0.032, *P* = 0.051; maize leaf prokaryotes R^2^ = 0.023, *P* = 0.418; soybean leaf fungi R^2^ = 0.018, P = 0.483; soybean leaf prokaryotes R^2^ = 0.035, *P* = 0.128). Despite this, changes to the fungal phyllosphere composition by fungicide treatments differed depending on management (fungicide-management interaction) only in soybean, and not in maize the phyllosphere (maize leaf fungi 9-dpf R^2^ = 0.012, *P* = 0.916; soybean leaf fungi 13-dpf R^2^ = 0.041, *P* = 0.017; soybean leaf fungi 33-dpf R^2^ = 0.039, *P* = 0.015). There was no substantial evidence that fungicides altered the composition of phyllosphere prokaryote communities, prokaryote root communities, or fungal root communities. Therefore, the variance explained due to the fungicide disturbance was examined for fungal phyllosphere communities before and after fungicide exposure while partitioning out the variation due to crop management by a constrained analysis of principal coordinates (CAP) (Fig. 1).

**Fig. 1.**
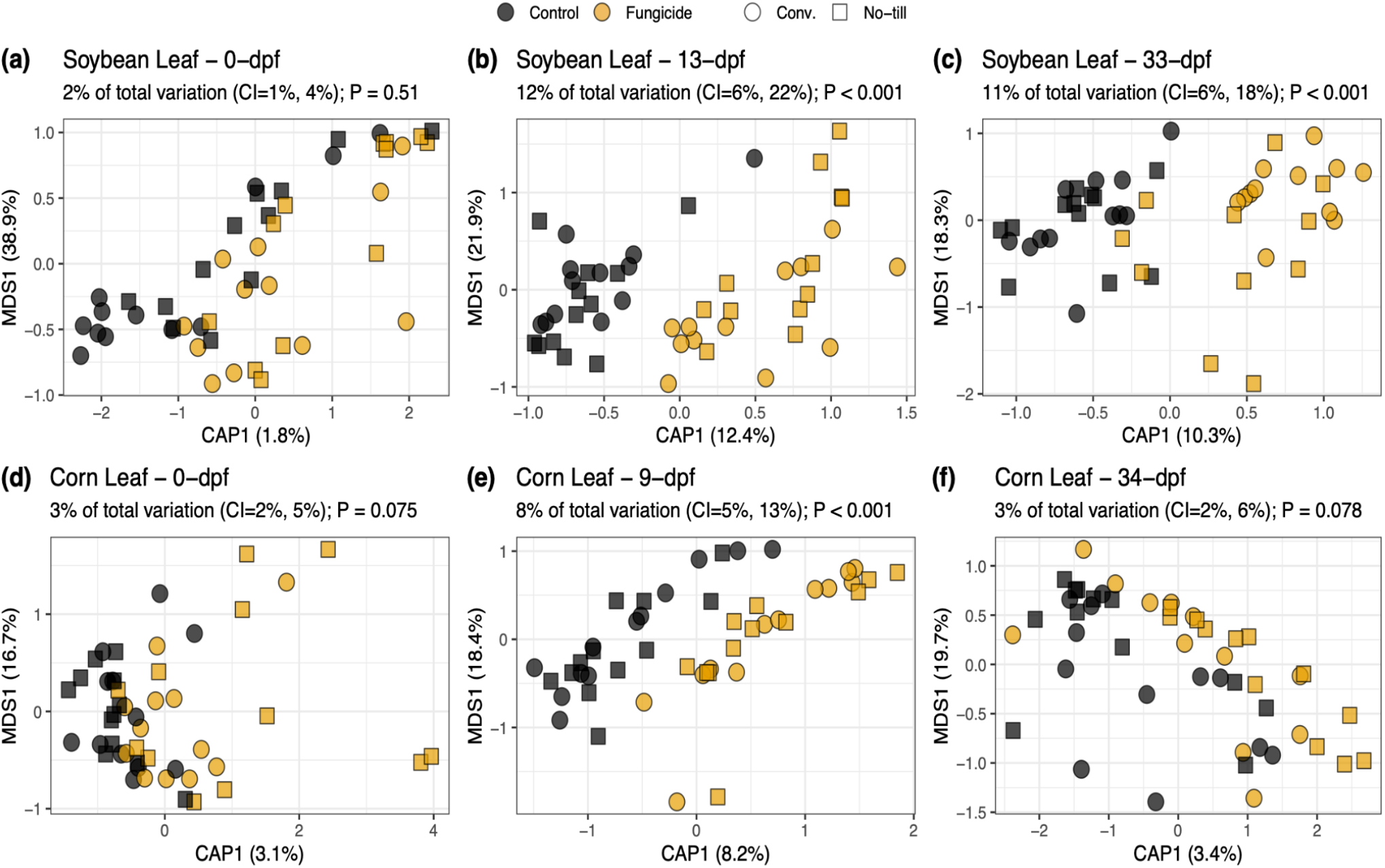
Effects of fungicides on fungal leaf composition in maize and soybean. Separate analyses were conducted for soybean (a) 0-(b) 13- or (c) 33-days post fungicide application or maize (d) 0-(e) 9- or (f) 34-days post fungicide application and days post fungicide application (dpf) since there was a significant interaction between dpf and fungicide. Constrained analysis of principal coordinates (CAP) analyses was constrained by the effect of fungicide while partialling out the effect of treatment. The percentage of total variation due to fungicide is expressed above the plot. The significance was determined based on 1000 permutations.

For soybean and maize leaves, no significant differences were observed prior to fungicide application (*P* = 0.51), but fungicide treatment had a significant effect on fungal leaf composition after fungicides were applied (13-dpf 13 % variation *P* < 0.001; 33-dpf 11 % variation *P* < 0.001) (Fig.1a-c). Similarly, the effect of fungicide disturbance on maize leaf fungal composition was not observed before fungicides were applied (*P* = 0.075) (Fig. 1d). However, unlike soybean, there was no evidence the fungicide altered fungal composition longer than nine days (9-dpf 8 % variation *P* < 0.001; 34-dpf 3 % variation *P* = 0.078) (Fig. 1e, 1f). The non-significant beta dispersion tests across managements at 9-dpf or 34-dpf for maize (9-dpf conventional *P* = 0.369, no-till *P* = 0.631; 34-dpf conventional *P* = 0.364, no-till *P* = 0.662) or 13- and 33-dpf (13-dpf conventional *P* = 0.742, conventional *P* = 0.866; 33-dpf conventional *P* = 0.335, no-till *P* = 0.123) for soybean, indicate that the effects of fungicide on fungal leaf composition are likely due to true differences in community composition rather than group dispersions (Table S6).

### Fungicidal effects on network properties depend on crop management

In soybean under conventional and no-till management, network complexity was not significantly different before fungicide application (conventional *P* = 0.13; no-till *P* = 0.93), but was significantly lower than control plots 13-dpf (conventional *P* = 0.0015; no-till *P* = 0.3) (Fig. 2a). However, after one month, the soybean no-till network complexity had recovered (*P* = 0.12), whereas the conventional treatment was still significantly lower compared to the non-sprayed control plots (*P* = 0.003) (Fig. 2a). The loss in network complexity can partially be explained by a reduction in the number of nodes (i.e., OTUs) since the average number of nodes per network also followed this same trend (Fig. 2b). These same effects were not observed in maize. Fungicide disturbance was followed by the loss of network complexity mainly through node loss, but crops and crop management impacted network properties under fungicide disturbance. To investigate these trends more closely, we investigated the specific fungal taxa affected through differential abundance analysis.

**Fig. 2.**
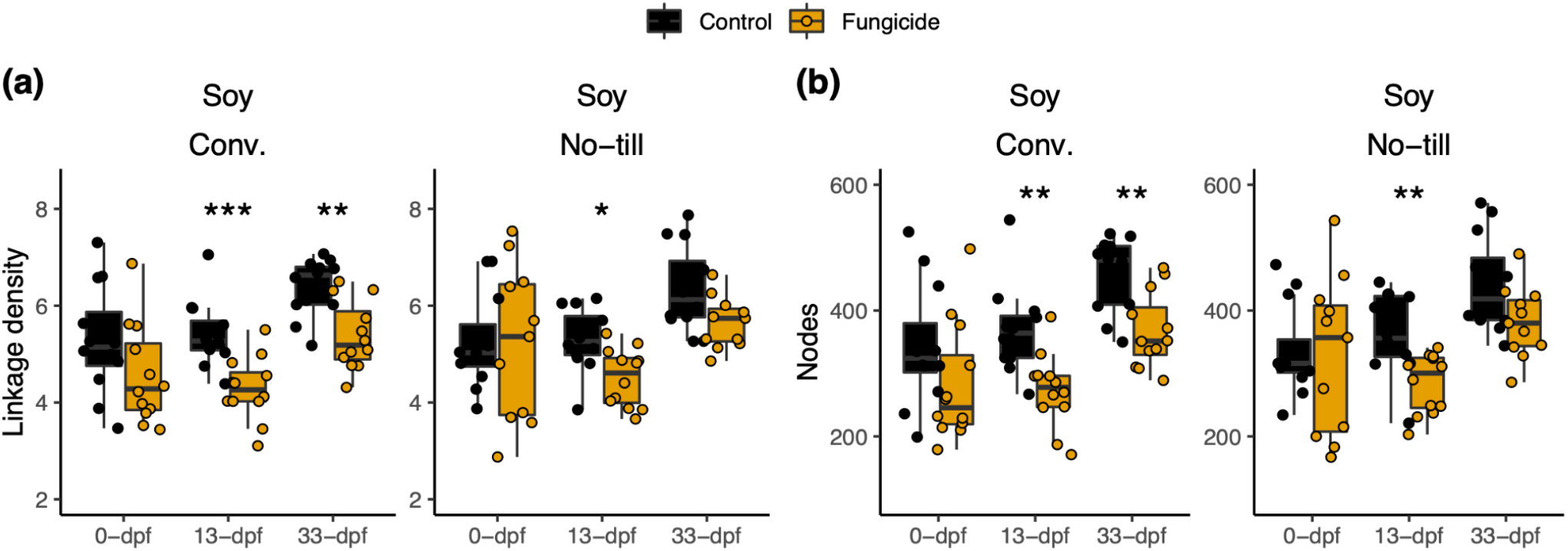
Fungicides alter soybean network complexity. A microbial co-occurrence network was constructed using taxa with a mean relative abundance greater than 1^-5^ and present in greater than 5 % of samples. Subnetworks were created for each sample based on the OTUs present, and each point represents a subnetwork. (a) Network complexity (i.e., linkage density) and (b) number of edges were then calculated for each subnetwork. Comparisons are based on Wilcox ranked sign tests for soybean conventional management and no-till. Asterisks indicate the level of significance; * = p ≤ 0.05, ** = p ≤ 0.01, *** = p ≤ 0.001

### Identification of fungicide-affected fOTUs

To determine which fungal taxa were significantly affected by fungicide application, a differential abundance analysis was conducted with ANCOM (Table S7). In total, the abundance of 238 unique fOTUs representing 21 fungal classes was altered by fungicide treatments across the two crops. Ascomycota (52.9%) and Basidiomycota (43.3%) fOTUs made up 96.2% of the differentially abundant fOTUs. Within Ascomycota, the Dothidiomycetes (28.6%) and Sordariomycetes (9.66%) accounted for the largest percentage of fOTUs that were differentially abundant following fungicide treatment (Fig. 3a). These fungi may be expected since many foliar plant pathogens fall within this class of fungi, and fungicides typically target these pathogen groups. Unexpectedly, a large percentage of fOTUs that were differentially abundant included non-target dimorphic clades of fungi that commonly exist as yeasts such as Agaricostilbomycetes, Cystobasidiomycetes, Exobasidiomycetes, Microbotryomycetes, Spiculogleomycetes, Taphrinomycetes, and Tremellomycetes. A total of 83 fOTUs across these classes were significantly different in abundance following the fungicide application in maize or soybean (Fig. 3a). Notably, Tremellomycetes made up the second-largest class of fungi (42 fOTUs, 17.6%) differentially abundant. Of the Tremellomycetes, 57.1% were concentrated within the Bulleribasidiaceae, accounting for 24 fOTUs that were differentially abundant compared to non-sprayed control. Twenty-three of the Bulleribasidiaceae significantly decreased in abundance.

**Fig. 3.**
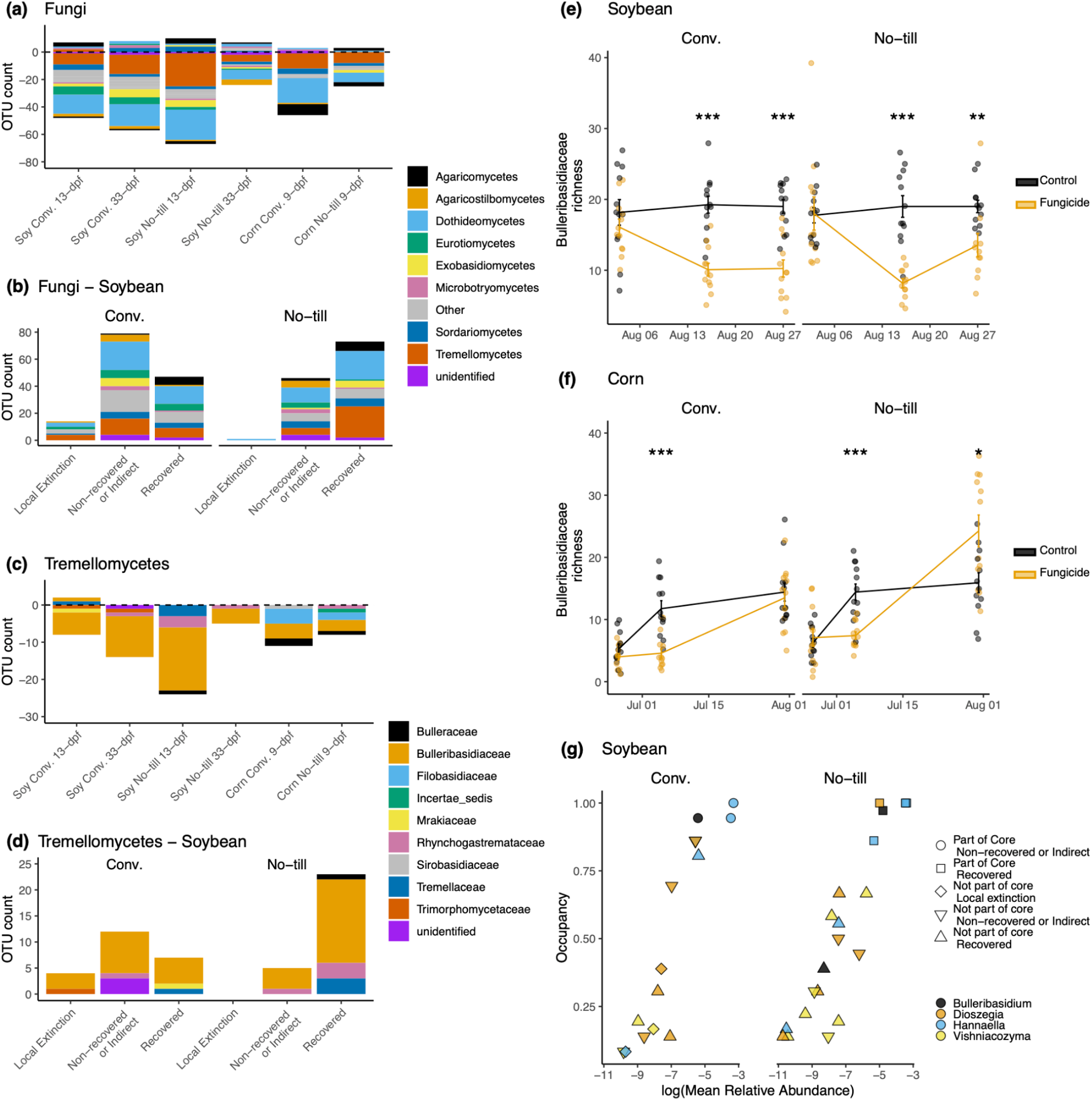
Management scheme alters the recovery dynamics of phyllosphere fungi following fungicide treatment. The composition of fungal operational taxonomic units (OTUs) significantly different in abundance, as indicated with analysis of compositions of microbiomes (ANCOM) analysis (n = 12). (a) Composition of fOTUs whose abundance was significantly different following a fungicide disturbance. Bars below zero indicate the fOTU decreased in abundance, whereas bars above zero indicate the fOTU increased in abundance. (b) Recovery of fungi in soybean leaf samples in conventional (conv.) or no-till management. (c) Composition of fOTUs within the Tremellomycetes whose abundance was significantly altered following a fungicide disturbance. (d) Recovery dynamics of Tremellomycetes fOTUs following a fungicide disturbance in conv. or no-till (e) soybean or (f) maize plots subjected to a fungicide treatment compared to non-sprayed control plots. (g) Abundance occupancy relationship with the recovery dynamics of the Bulleribasidiaceae fOTUs significantly affected by the fungicide treatment. Asterisks indicate the level of significance; * = p ≤ 0.05, ** = p ≤ 0.01, *** = p ≤ 0.001

However, not all yeast fOTUs decreased in abundance. For example, *Bulleromyces albus* fOTU10 increased in relative abundance 4.25 times in soybean conventional management 13-dpf but was not significantly different than the control after 33-dpf. In contrast, two *Sporobolomyces* fOTUs (fOTU66 and fOTU94) increased relative abundance following fungicide application in soybean and remained significantly (7 times) higher in fungicide treated plots than in control plots 33-dpf. *Sporobolomyces patagonicus* fOTU94 was 4.38 times more abundant in the fungicide treated plots than the control 13-dpf in the conventional management and remained significantly higher in fungicide sprayed plots (9.06 times) compared to the control after 33-dpf. *Sporobolomyces roseus* fOTU66 was 15 times more abundant in the conventionally managed fungicide treated plots 33-dpf. This same increase in *Sporobolomyces* abundance was not observed in maize.

### Resilience of the core mycobiome and local extinctions of accessory members

Many of the fOTUs affected by a fungicide application were also part of the core phyllosphere taxa (Table S8). In conventionally managed soybean plots, 22 fOTUs were determined to be core fungal phyllosphere taxa, and the abundances of five of these core members (fOTU 6 *Mycosphaerella* sp., fOTU 10 Tremellales, fOTU 34 *Hannaella* sp., fOTU 13 *Hannaella* sp., and fOTU 643 *Tilletiopsis* sp.) were significantly different following fungicide application. *Hannaella* sp. (fOTU 13), *Hannaella* sp. (fOTU 13), and *Tilletiopsis* sp. (fOTU 643) were also part of the 43 core members of the no-till soybean phyllosphere affected by fungicide application. Of the 40 core members of the conventionally managed maize phyllosphere, the abundance of four Tremellomycetes fOTUs and one unidentified fungal taxon (fOTU 116) were significantly different following fungicide application. These included three yeast taxa that were not members of the soybean core, which included two fOTUs in the genus *Filobasidium* (fOTU 82 *Filobasidium oeirense*, and fOTU 97 *Filobasidium*), one *Bullera crocea* (fOTU 65), and *Vishniacozyma globispora* (fOTU 83). Two of these fOTUs (fOTU 97 *Filobasidium* and fOTU 65 *Bullera crocea*) were also core members of the maize phyllosphere in the no-till management that were significantly altered by the fungicide.

None of the core members of the phyllosphere taxa became locally extinct following fungicide application in the core microbiome of both crop managements. However, among the taxa whose abundance was significantly altered by the fungicide application in soybean, the no-till management had a 61 % recovery compared to the 34 % recovery of the conventionally managed soybean (Fig. 3b; Table S9). Fourteen fungal OTUs became locally extinct following fungicide application in soybean managed conventionally compared to one in the no-till plots (Fig. 3b). Among the Tremellomycetes fOTUS whose abundances were significantly impacted by fungicide applications, the majority were Bulleribasidiaceae (Fig. 3c). Eighty-two percent of affected Bulleribasidiaceae recovered in no-till managed soybean compared to the 30% of the conventionally managed plots. No Bulleribasidiaceae taxa became locally extinct in the no-till plots; in contrast, three Bulleribasidiaceae fOTUs were never observed following fungicide application in the conventional management (Figure 3d). The trend of increased recovery was also evident in Bulleribasidiaceae richness on the last sampling for maize (33-dpf) and soybean (34-dpf) no-till samples (Fig. 3e, 3f). Bulleribasidiaceae in the core of conventional management did not fully recover within the study period (Fig. 3g). Additionally, the Bulleribasidiaceae in the conventional management that were locally extinct following fungicide disturbance occupied less than 50% samples in non-sprayed plots indicating that local extinctions caused by fungicides affect the rare, non-core members of the community (Fig. 3g). No local extinctions among fungal taxa were detected in maize fungicide treated plots; all impacted taxa recovered.

### Indirect effects of fungicides on prokaryotes mediated through yeast

Random forest models based on prokaryotic abundance on soybean leaves sprayed with fungicides explained a significant amount of variance (*P* < 0.001) in Bulleribacideaceae richness in the no-till (28.70%; R^2^ = 0.25) and conventional (43.47%; R^2^ = 0.44) management (Fig. 4a, 4b). We then extracted the set of OTUs that were the most important for maintaining the model’s accuracy in fungicide treated plots. However, there was no evidence (*P* ≥ 0.05) those same taxa were able to predict Bulleribacideaceae richness in control samples indicating the unique effect of the fungicide (Fig. S3). OTUs classified as *Sphingomonas, Methylobacterium*, and *Hymenobacter* were the most important for predicting fungal richness in the no-till management (Fig. 4a, 4b). Many taxa from the same genera were important in predicting Bulleribasidiaceae richness in the conventional management system, including the *Sphingomonas* and *Hymenobacter* genera (Fig. 4c, 4d). However, other genera were unique by management type, including *Methylobacterium* for the no-till management system and *Pseudokineococcus* and *Kineococcus* in the conventional management system

**Fig. 4.**
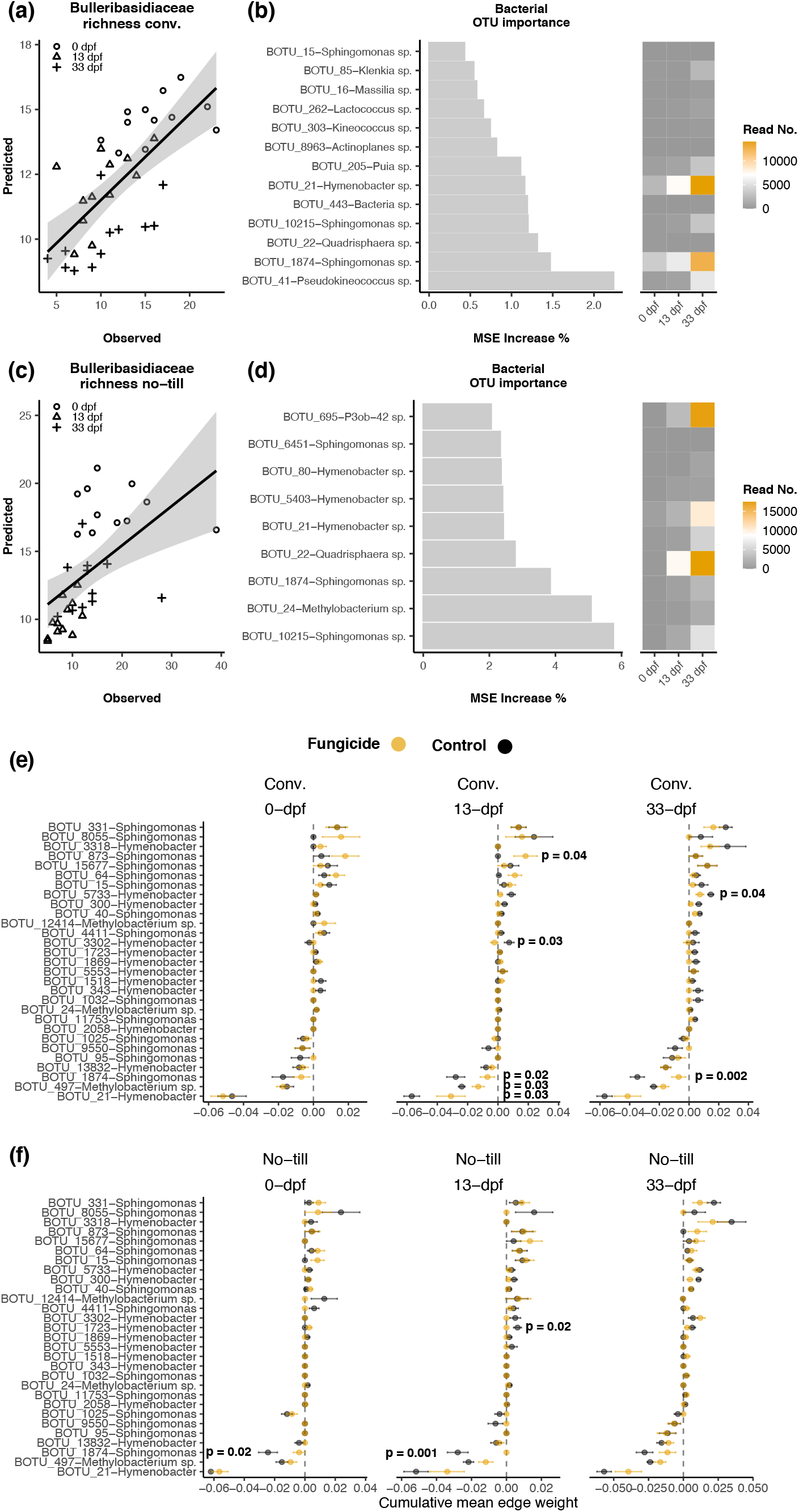
Indirect effects of fungicides on prokaryotic communities from altered Bulleribacideaceae diversity on soybean leaves. Relationship of observed versus predicted Bulleribasidiaceae richness in conventional (a) or no-till (c) from random forest models using prokaryote OTU abundance in fungicide treated plots. The most important (*P* < 0.05) prokaryote OTUs for random forest model accuracy in fungicide treated conventional (b) or no-till (d). The cumulative mean edge weight calculated from each sub-network of a meta-network of Bulleribacidaceae edges between *Sphinogomonas, Hymenobacter*, or *Methylobacterium* OTUs and alterations to co-occurrence strength with and without fungicides under (e) conventional and (f) no-till crop management.

Prokaryote OTUs that were important for random forest model accuracy increased in abundance in fungicide treated plots as a response to altering Bulleribacideaceae diversity and were negatively co-associated with Bulleribacideaceae (Fig. 4b, 4d, 4e, 4f). For example, pOTU21 *Hymenobacter* and pOTU1874 *Sphingomonas* abundance increased as Bulleribacideaceae richness decreased and was negatively co-associated with the Bulleribacideaceae (Fig. 4b, 4e, 4f). Additionally, the cumulative edge weight between pOTU21 *Hymenobacter*, pOTU1874 *Sphingomonas*, and Bulleribacideaceae significantly changed when sprayed with fungicides in the conventional management, but not always in the no-till management, indicating that a loss in Bulleribacideace diversity can indirectly influence the co-occurrence between fungi and bacteria in different crop management schemes. However, not all co-occurrences between the Bulleribacideaceae and prokaryotes were negative, indicating that positive co-occurrences between prokaryotes and fungi in the phyllosphere may shift as well (Fig. 4e, 4f).

## Discussion

To our knowledge, this is the first study to assess the effect of fungicide-imposed disturbance and resiliency under different agricultural management systems. We found that fungicide applications had little direct effect on root fungal or prokaryotic communities but a substantial effect on target and non-target fungal phyllosphere communities and minor indirect effects on prokaryote communities in the phyllosphere. Soil fungi and prokaryotes were also sequenced in soybean, and there was no evidence of fungicidal effects (data not shown). Leveraging the KBS LTER site allowed the direct comparison of long-term crop management impacts to the microbiome without confounding location. Our data demonstrate that the phyllosphere microbiome’s resilience depends on the cropping management system, with a greater recovery in the abundance of affected phyllosphere microbiota in long-term no-till compared to annually tilled conventional management. Among the most important results was the commonality in the fungal taxa affected by fungicide treatments, particularly non-targeted groups of phyllosphere yeasts across crops. In maize and soybean, fungi in Dothidiomycetes (target) and Tremellomycetes (non-target) decreased in abundance following fungicide applications, raising questions on the role of Tremellomycete yeasts; specifically, the Bulleribasidiaceae play in phyllosphere microbiomes, and the effects of fungicide used in the absence of disease pressure.

This study observed reductions or local extinctions of these yeasts following fungicide application, which may lead to unintended consequences in the host plant. Phyllosphere yeast communities have received less attention in the literature than prokaryotes (Lindow & Brandl, 2003). The three Bulleribacideaceae genera observed in this study were *Hannaella, Dioszengia*, and *Vishniacozyma, Dioszengia*, and *Hannaella* have been demonstrated to produce the plant growth-promoting hormone indole acetic acid (IAA), similar to many plant growth-promoting phyllosphere prokaryotes (Sun *et al*., 2014; Wang *et al*., 2016). In comparison, *Vishniacozyma* yeasts have remained understudied but have been isolated from maize kernels (Yurkov & Kurtzman, 2019). Agler *et al*. (2016) and Wang *et al*. (2016) found *Dioszegia* was a hub taxon important in maintaining fungal-prokaryote interactions by altering prokaryote diversity in the phyllosphere microbiome of *Arabidopsis*. Importantly, in the absence of disease pressure as observed in this study, our results indicate that fungicide applications may affect populations of resident beneficial microbes. However, adverse impacts would be expected to be outweighed if the fungicide mitigates the disease.

Here, we show for the first time that fungicidal impacts on crop microbiomes is dependent on crop management, addressing a knowledge gap that previous studies were unable to address specifically (Karlsson *et al*., 2014; Sapkota *et al*., 2015; Knorr *et al*., 2019). A higher proportion of fOTUs altered by fungicide application in the no-till management system showed improved resilience within the study period, which may be explained by the differences in microbial communities present in the phyllosphere of each management before fungicide applications, as has been demonstrated previously at the KBS LTER site (Longley *et al*., 2020; Gdanetz & Trail, 2017). A previous study from the KBS LTER site has demonstrated that aerially dispersed yeasts are enriched in the phyllosphere but also found in lower abundance in belowground plant organs (Gdanetz *et al*., 2021). Crop residue left from previous seasons can harbor fungi that potentially may act as a source to repopulate the phyllosphere following a disturbance like the phenomenon of pathogens transferring from residues (Sommermann *et al*., 2018). Yeasts that inhabit the phyllosphere are primarily known to disperse through ballistosporic aerial dispersal, and the reassembly of leaves following fungicides may rely heavily on this spore dispersal mechanism. However, not all yeast taxa in the Bulleribacideaceae have been observed to form ballistocondia in culture (Li *et al*., 2020), leaving arguably less efficient means of dispersal from insects or through wind and rain (Gilbert, 1980; Starmer *et al*., 1988). Regardless, locally extinct taxa were not part of the core microbiome regardless of management systems or spore dispersal mechanism, demonstrating a tight relationship between abundance-occupancy and disturbance. These results demonstrate that microbiome resilience is improved in no-till crop management, informs discussion on managing crops for resilience, and demonstrates an ecosystem service provided by no-till agriculture in addition to improved nutrient cycling or preservation of habitats for microorganisms and mesofauna (Murrell, 2017).

In addition to differential impacts by crop management, fungicide applications affected soybean and maize phyllosphere communities differently. These differences may be due to crop, planting year, or fungicide regime. The effect of fungicide was likely reduced in the final sampling of maize due to sampling new leaves not directly sprayed, indicating that any effect would have been through the systemic activity of the fungicide 34 days after, which may have decreased the effect on the microbiome. Another critical difference is that the Delaro^®^ fungicide applied to soybeans in 2018 has two modes of action. Application of fungicides having two different modes of action has been shown to have a more significant effect on fungal beta diversity than a single mode of action in cereal crops (Sapkota *et al*., 2015). Although the impact of fungicides varied in magnitude between the two crops, the commonality of off-target impacted taxa between crops and fungicides demonstrates that multiple fungicide products on different crops consistently reduce these taxa. This information can be used to inform decisions on the use of fungicides under low pathogen pressure across multiple crops and cropping systems.

Recovery of network complexity is one measure of microbiome resilience. We show that network complexity decreased significantly in the soybean phyllosphere following fungicide treatment. Other studies have demonstrated that agricultural management alters network complexity. However, the functional consequences of these changes were not directly assessed (Banerjee *et al*., 2019; Schmidt *et al*., 2019). In soils, it has been demonstrated that increases in network complexity are positively correlated with various ecosystem functions and increases in the number of unique functions and functional redundancy (Wagg *et al*., 2019). The functional consequences of decreases in network complexity remain unexplored in the phyllosphere microbiome. They may provide the rationale for chemical application decisions or novel microbial-based treatments to replace lost taxa.

Notably, fungicide application altered co-occurrences between phyllosphere fungi and prokaryotes, demonstrating the indirect effects of fungicide applications through the loss in the diversity of Bulleribacideaceae. In support of random forest results, many of the same prokaryotes identified from networks as having changes in cumulative mean edge weight were identified by random forest as predicting Bulleribasidiaceae richness. Disturbance can change cooperation/competition dynamics, and a high level of disturbance can reduce cooperation (Brockhurst *et al*., 2007; 2010). In our study, the cumulative mean edge weight between most phyllosphere prokaryotes and Bulleribasidiaceae became more positive, indicating fewer negative associations between a particular bacterium and the Bulleribasidiaceae. However, there were exceptions where cumulative edge weights, positive before spray became neutral, likely due to the disappearance of some fungal taxa from samples following fungicide application, and therefore the disappearance of any associations. However, loss of negative correlations may also be due to reduced competition between phyllosphere prokaryotes and Bulleribasidiaceae as more niche space is available to phyllosphere prokaryotes following fungicide application.

Shifts in correlations between Bulleribasidiaceae and phyllosphere prokaryotes are of interest due to the unique physiology of many phyllosphere prokaryotes as it relates to plant health. *Methylobacterium* spp. have been demonstrated to be abundant in plants’ phyllosphere and have the genes to produce plant growth-promoting auxins and UVA-absorbing compounds (Kwak *et al*., 2014; Yoshida *et al*., 2017). *Hymenobacter* sp., *Methylobacterium* sp., and *Sphingomonas* sp. are core phyllosphere members in switchgrass (Grady *et al*., 2019) and are highly abundant in the *Arabidopsis* phyllosphere (Delmotte *et al*., 2009).

A comprehensive view of the phyllosphere organisms is needed to understand microbiome functioning and plant health. This research demonstrates that foliar fungicide treatments alter phyllosphere microbiomes in maize and soybean, and non-target Bulleribacideaceae yeasts were negatively impacted in soybean and maize phyllospheres. Microbiome complexity was altered partially by decreasing co-occurrence between Bulleribacideaceae yeasts and dominant phyllosphere prokaryote taxa, demonstrating indirect effects of fungicide applications mediated through the presence of these yeasts. Further, these data support our hypothesis that the recovery of the phyllosphere microbiome differed by tilling management. Together, these results improve our understanding of fungicide impacts on crop microbiomes and their recovery in different managements and inform their rational use to maintain efficacy and intended impacts across different cropping systems.

## Acknowledgments

This work was supported by the NIFA grant MICL08541 from the USDA National Institute of Food and Agriculture to GB, FT, and MC and USDA MICL02416 to GB. Support for this research was also provided by the NSF Long-term Ecological Research Program (DEB 1832042) at the Kellogg Biological Station and by Michigan State University AgBioResearch. We would also like to thank Elizabeth Gall for field and lab assistance. We would like to thank Alex Witte for help with applying the fungicides.

## Author Contributions

ZAN and RL contributed equally to this work. MIC, FT, and GB designed experiments. MIC applied fungicides. MIC, FT, GB, and RL collected samples. ZAN and RL prepared amplicon libraries and generated sequencing data. ZAN, RL, GMNB, and GB wrote and edited the manuscript, with contributions from MIC and FT. ZAN, RL, GMNB processed sequencing data and analyzed data. ZAN generated figures.

### Data availability

Raw sequences for soybean microbial communities used to create figures and tables in this study are available in the NCBI SRA database under the following accession numbers: PRJNA603199 and PRJNA603207. Sequences which were produced on the same Miseq runs and used to remove contaminants are available in PRJNA603147. Raw sequences for maize microbial communities are available under the following accession numbers: PRJNA739465 and PRJNA739759. Code to generate figures and tables are located on GitHub at https://github.com/noelzach/FungicidePulseDisturbance

## Figures and Tables

**Fig. 1 Effects of fungicides on fungal leaf composition in maize and soybean**. A separate analysis was conducted for soybean (a) 0-(b) 13- or (c) 33-days post fungicide application or maize (d) 0-(e) 9- or (f) 34-days post fungicide application and days post fungicide application (dpf) since there was a significant interaction between dpf and fungicide. Constrained analysis of principal coordinates (CAP) analyses was constrained by the effect of fungicide while partialling out the effect of treatment. The percentage of total variation due to fungicide is expressed above the plot. The significance was determined based on 1000 permutations.

**Fig. 2 Fungicides alter soybean network complexity**. A microbial co-occurrence network was constructed using taxa with a mean relative abundance greater than 1^-5^ and present in greater than 5 % of samples. Subnetworks were generated for each sample based on the OTUs present within those samples, and each point represents a subnetwork. (a) Network complexity (i.e., linkage density) and (b) number of edges were then calculated for each subnetwork. Comparisons are based on Wilcox ranked sign tests for soybean conventional management and no-till. Asterisks indicate the level of significance; * = p ≤ 0.05, ** = p ≤ 0.01, *** = p ≤ 0.001

**Fig. 3 Management scheme alters the recovery dynamics of phyllosphere fungi following fungicide treatment**. The composition of fungal operational taxonomic units (OTUs) significantly different in abundance, as indicated with analysis of compositions of microbiomes (ANCOM) analysis (n = 12). (a) Composition of fOTUs whose abundance was significantly different following a fungicide disturbance. Bars below zero indicate the fOTU decreased in abundance, whereas bars above zero indicate the fOTU increased in abundance. (b) Recovery of fungi in soybean leaf samples in conventional (conv.) or no-till management. (c) Composition of fOTUs within the Tremellomycetes whose abundance was significantly altered following a fungicide disturbance. (d) Recovery dynamics of Tremellomycetes fOTUs following a fungicide disturbance in conv. or no-till (e) soybean or (f) maize plots subjected to a fungicide treatment compared to non-sprayed control plots. (g) Abundance occupancy relationship with the recovery dynamics of the Bulleribasidiaceae fOTUs significantly affected by the fungicide treatment. Asterisks indicate the level of significance; * = p ≤ 0.05, ** = p ≤ 0.01, *** = p ≤ 0.001

**Fig. 4 Indirect effects of fungicides on prokaryotic communities from altered Bulleribacideaceae diversity on soybean leaves**. Relationship of observed versus predicted Bulleribasidiaceae richness in conventional (a) or no-till (c) from random forest models using prokaryote OTU abundance in fungicide treated plots. The most important (*P* < 0.05) prokaryote OTUs for random forest model accuracy in fungicide treated conventional (b) or no-till (d). The cumulative mean edge weight calculated from each sub-network of a meta-network of Bulleribacidaceae edges between *Sphinogomonas, Hymenobacter*, or *Methylobacterium* OTUs and alterations to co-occurrence strength with and without fungicides under (e) conventional and (f) no-till crop management.

## Supplemental Information

**Fig. S1**. Rarefaction curves for each sample sequenced in this study for (a) fungi and (b) prokaryotes in soybean or maize leaves and roots. The dashed line represents the median sequence depth.

**Fig. S2**. Fungicidal effects on the richness of different fungal classes in soybean and maize phyllosphere. Black dots are control yellow dots are fungicide samples. Asterisks indicate the level of significance; * = p ≤ 0.05, ** = p ≤ 0.01, *** = p ≤ 0.001

**Fig. S3**. Random forest models percent explained variance, error, and overall model significance (permutations = 999) for (a) conventional management treated with fungicides, (b) no-till treated with fungicides, (c) conventional management control, and (d) no-till control.

**Table S1**. Three-step amplicon library preparation reagents and PCR master mixes

**Table S2**. Primers used for amplicon library preparation

**Table S3**. Cycling conditions for PCR in library preparation for fungi and prokaryotes

**Table S4**. Permutational multivariate analysis of variance for fungi in maize or soybean in roots or leaves before and after fungicide application

**Table S5**. Permutational multivariate analysis of variance for prokaryotes in maize or soybean in roots or leaves before and after fungicide application

**Table S6**. Effects of fungicide on maize and soybean leaf fungal composition

**Table S7**. Differentially abundant phyllosphere fungal OTUs by fungicide treatment

**Table S8**. Core members of the soybean or maize phyllosphere in no-till and conventional management

**Table S9**. Recovery status of fungicide-impacted soybean phyllosphere fungal OTUs

