## Supplementary figures and images for "Non-target impacts of fungicide disturbance on phyllosphere yeasts in different crop species and management systems"

### FigS1

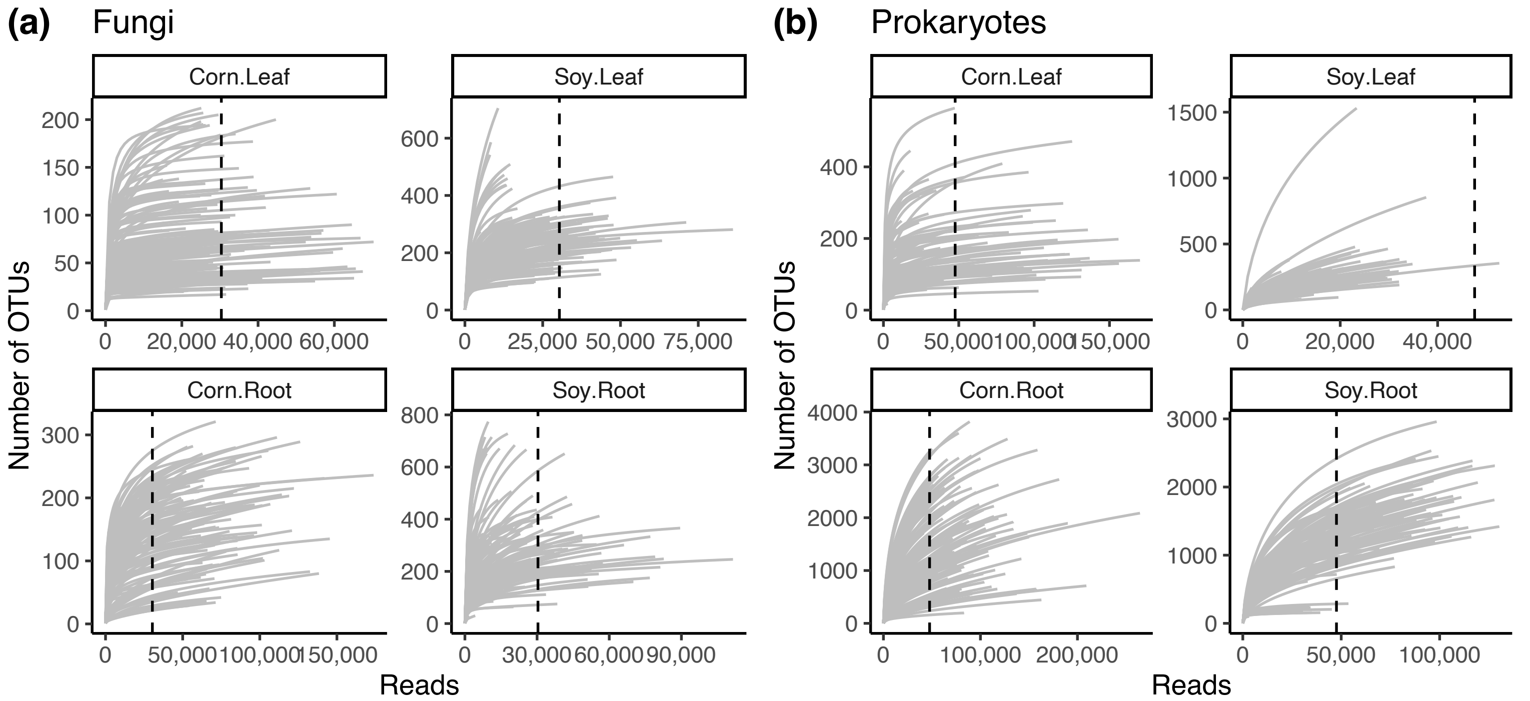

### FigS2

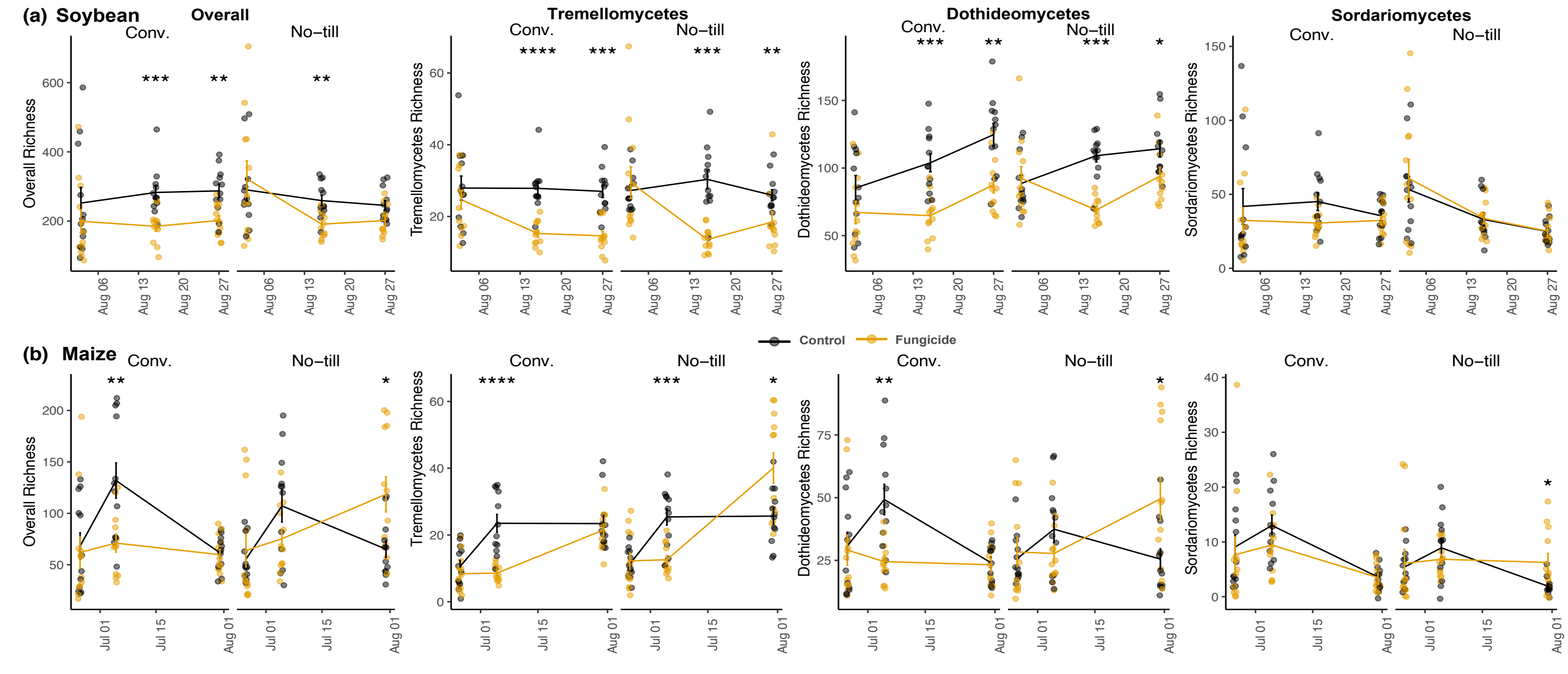

### FigS3

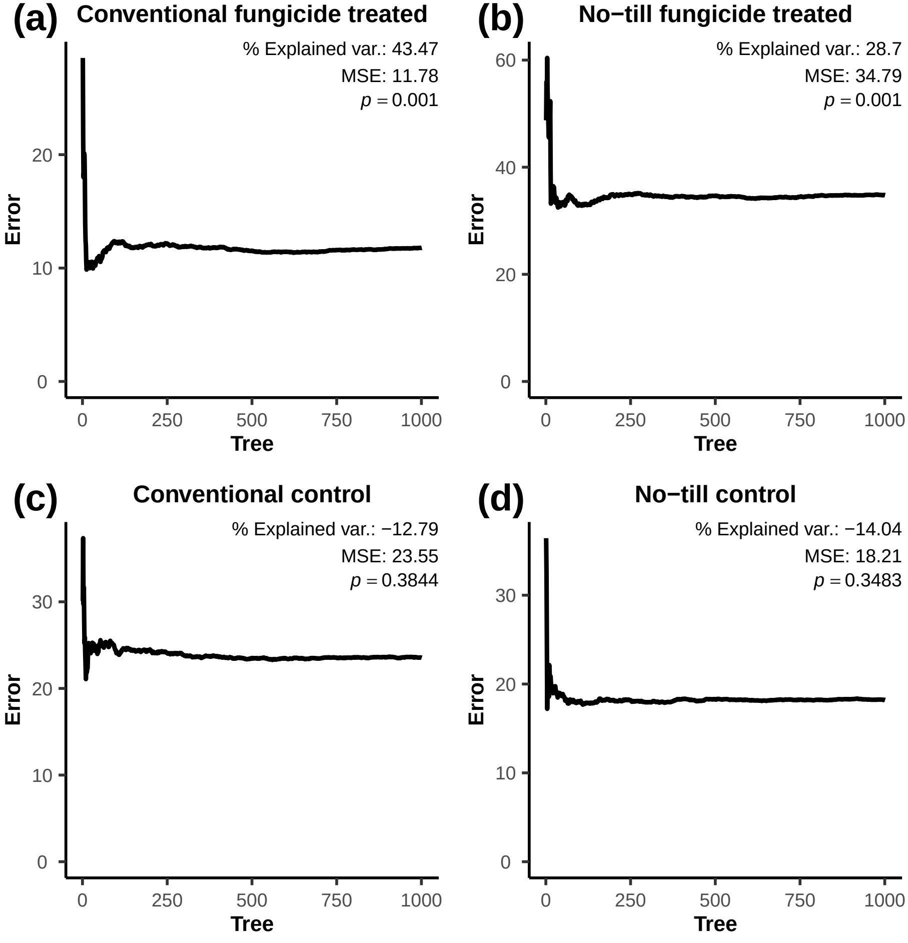
